# DropSynth 2.0: high-fidelity multiplexed gene synthesis in emulsions

**DOI:** 10.1101/740977

**Authors:** Angus M. Sidore, Calin Plesa, Joyce A. Samson, Sriram Kosuri

## Abstract

Multiplexed assays allow functional testing of large synthetic libraries of genetic elements, but are limited by the designability, length, fidelity and scale of the input DNA. Here we improve DropSynth, a low-cost, multiplexed method which builds gene libraries by compartmentalizing and assembling microarray-derived oligos in vortexed emulsions. By optimizing enzyme choice, adding enzymatic error correction, and increasing scale, we show that DropSynth can build thousands of gene-length fragments at >20% fidelity.

## Main Text

Multiplexed functional assays link gene function or regulation to activities that can be read by next-generation sequencing such as through enrichment screens (cellular growth^1^, cell sorting^2,3^, binding^4,5^) or transcriptional reporters^6^. Multiplexed assays can functionally assess thousands of different sequences in a single pooled experiment, and are thus powerful approaches for understanding how sequence affects function^7^. The DNA sequences to test are produced by genome fragmentation^8^, mutagenesis of existing sequences^9^, or by direct synthesis of oligonucleotides (oligos)^10^. Direct oligo synthesis allows for testing controlled hypotheses against one another without the constraints of natural variation or mutagenesis. However, individual oligos are generally shorter than 200 nucleotides (nt), limiting potential applications. Gene synthesis from oligo libraries can be used to extend these lengths^11,12^, but the high cost of individual assembly and processing becomes prohibitive for large gene libraries.

To address these concerns, we previously developed a low-cost, multiplexed method termed DropSynth, which is capable of building large gene libraries from microarray-derived oligos^13^. DropSynth works by assembling genes through the isolation and assembly of microarray-derived oligos in droplets (Fig. 1a). First, genes are bioinformatically split into several oligos and flanked with restriction sites, priming sequences, and a 12 nt microbead barcode sequence that is common to all oligos needed to assemble a given gene (Supplementary Fig. 1). Oligos are synthesized as a microarray-derived pool, amplified and nicked using a nicking endonuclease, exposing each 12 nt microbead barcode as a single-stranded overhang. Nicked oligos are hybridized to a pool of barcoded microbeads that contain complementary 12 nt microbead barcode sequences, such that each bead pulls down all oligos for a particular assembly. Bound beads are then encapsulated in droplets, where sequences are cleaved from the bead using a IIS restriction enzyme and assembled into genes using a high fidelity polymerase. Following assembly, the emulsion is broken and gene libraries are recovered.

**Fig. 1.**
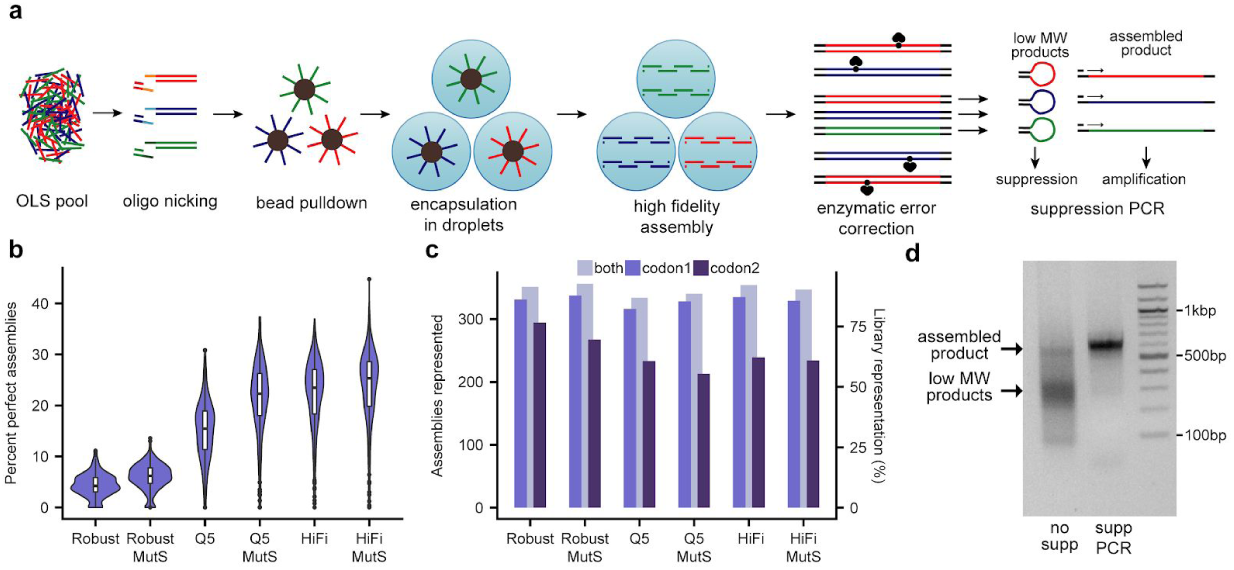
DropSynth 2.0: high-fidelity multiplexed gene synthesis in emulsions. **a**, Schematic of DropSynth 2.0. **b**, Comparison of percent perfect assemblies (minimum 100 assembly barcodes) of a 384-gene library assembled using DropSynth with 3 different polymerases (KAPA Robust, NEB Q5, or KAPA HiFi) with or without MutS-based enzymatic error correction. **c**, Comparison of total assemblies represented with at least one assembly barcode for all conditions. 2 codon versions of the 384-gene library were assembled for each condition, and representation is improved when combining across both codon usages. **d**, 2% agarose gel of 384-gene assembly product following bulk amplification with standard PCR or using single-primer suppression PCR; yield of assembled product is noticeably higher using single-primer suppression PCR.

However, DropSynth is limited by the resulting fidelity of the gene libraries and the scalability of the method. For example, in our original work, only 1.9-3.9% of assemblies corresponded to the designed protein sequence, and each assembly was limited to 384 designs per library^13^. This error rate and small scale inhibit broader applicability, but also limit its use as a broader gene synthesis method. Here we present DropSynth 2.0, an optimized protocol for multiplexed gene synthesis. We optimized enzyme choice, oligo design, assembly protocols, added enzymatic error correction, and increased scale, which together result in a substantially superior method for gene library synthesis.

The error profiles of previous DropSynth assemblies had many more transition mismatches than single-base deletions, the dominant error type of the oligos. This mutational signature was indicative of errors introduced by KAPA Robust polymerase, which was initially chosen for assembly performance^13,14^. We thus optimized two high-fidelity polymerases, KAPA HiFi and NEB Q5, with order-of-magnitude lower error rates^15^ to work with DropSynth. We assembled two codon versions of a 384-member library of dihydrofolate reductase (DHFR) homologs using KAPA Robust and the two high-fidelity polymerases. We found the high-fidelity polymerases produced less assembled product, and thus made cloning and size-selection difficult. To address this, we amplified the resultant assemblies using single-primer suppression PCR. In this technique, primer annealing competes with the self-annealing of inverted terminal repeats (ITRs) flanking the assembled genes^16,17^. Shorter by-products tend to self-anneal, while correct assembly products anneal to the primer, resulting in proper amplification. Following suppression PCR, we ligated the libraries into a plasmid containing a 20 basepair (bp) assembly barcode sequence, cloned and sequenced them, allowing us to link assembled genes with unique barcodes.

Amongst genes with at least 100 assembly barcodes, we found a median of 4.2% perfect assemblies at the amino acid level for KAPA Robust (Fig. 1b), which is consistent with our previous work^13^. Using high-fidelity polymerases for assembly resulted in a several-fold improvement in the median percent perfect assemblies, with 15.5% using NEB Q5 and 23.5% using KAPA HiFi. A similar trend in percent perfect assemblies was observed from the 2nd codon version assembled (Supplementary Fig. 2). When analyzing the total number of constructs represented with at least 1 assembly barcode, we found consistently high representation across all polymerases (86% for KAPA Robust, 87% for KAPA HiFi, and 82% for NEB Q5) for codon 1 (Fig. 1c). Codon 2 had lower library representation, particularly for NEB Q5 and KAPA HiFi (77% for KAPA Robust, 62% for KAPA HiFi, 61% for NEB Q5). Though differences in coverage exist between codon usages, combining across codon usages improves the total protein library representation (91% for KAPA Robust, 92% for KAPA HiFi, and 87% for NEB Q5) (Fig. 1c). Thus, by using multiple codon usages per gene, we improve our ability to achieve greater library coverage. Finally, we observed that using single-primer suppression PCR after assembly significantly improved the quantity of the correctly assembled product, while minimizing the presence of lower molecular weight by-products (Fig. 1d).

**Fig. 2.**
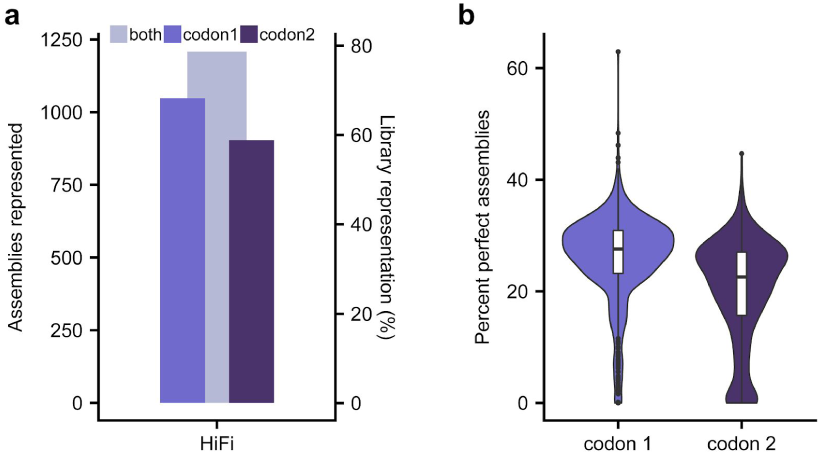
A scaled-up barcoded bead pool allows for the one-pot assembly of up to 1536 genes. **a**, 2 codon versions of a 1536-gene library were assembled using KAPA HiFi; when combining across both codon usages, 1208/1536 genes have at least one assembly barcode. **b**, Comparison of percent perfect assemblies (minimum 100 assembly barcodes) of both codon versions of each 1536-gene library.

We next set out to improve library representation by optimizing the algorithms that determine how oligos are split. Several factors can contribute to incomplete library representation, including oligo synthesis failure, processing failure, and assembly failure. One cause of assembly failure is the inability of oligos to overlap and assemble properly. In order to investigate this further, we created multiple iterations of the same two codon versions of our 384-member DHFR library using different parameters for where oligos overlapped, including overlap length and secondary structure^18^. We found that 20 bp overlaps had higher library representation than 25 bp overlaps, while modifying the overlap location to minimize secondary structure had minimal effect (Supplementary Fig. 3).

Assembly failure can also be attributed to incompatibilities between the polymerase buffer and the IIS restriction enzyme used to cleave oligos off the beads. In particular, NEB Q5 buffer inhibits several IIS restriction enzymes^19^, which can cause incomplete library representation by preventing the cleavage of oligos from the surface of the microbead within the droplet (Fig. 1c). To investigate this further, we designed multiple iterations of the same two codon usages of our 384-member DHFR library with three different IIS restriction sites (BtsI, BsmAI and BsrDI) and assembled them using NEB Q5. Though differences in library representation exist across codon versions, we found that assemblies using BsrDI had poor representation when compared to assemblies with BtsI and BsmAI. (Supplementary Fig. 4).

Experimental improvements to the workflow allowed us to test enzymatic mismatch correction techniques. Genes possessing mismatches or single-base insertions or deletions contain heteroduplexes after emulsion assembly, which can then be recognized and bound by the bacterial enzyme MutS^20,21^. Magnetic beads containing immobilized MutS capture these sequences, thus allowing for the enrichment of perfect genes. We found that using MutS-based error correction resulted in marginal improvements in fidelity (+2.0% for KAPA Robust, +6.8% for NEB Q5, +1.8% for KAPA HiFi) (Fig. 1b).

Finally, the scale per DropSynth assembly reaction was limited to 384 genes. In an effort to overcome this limitation, we designed and created a new barcoded bead pool containing 1536 unique microbead barcode sequences. This bead pool was constructed using similar procedures to the 384-plex bead pool (Supplementary Fig. 5, Supplementary Protocols). In order to demonstrate the efficacy of the new bead pool, we designed and assembled 2 codon versions of a 1536-member library of DHFR homologs. Each library member contains one of 1536 unique microbead barcode sequences which can be hybridized to one of 1536 beads with complementary barcode sequences. We assembled these libraries using our best-performing polymerase, KAPA HiFi, and ligated them into a barcoded expression plasmid. Following sequencing, we observed 1048/1536 (codon 1) and 904/1536 (codon 2) constructs represented with at least one assembly barcode (Fig. 2a). When combining across codon usages, we found a total of 1208 constructs represented, approaching 80% total protein library coverage (Fig. 2a). Amongst genes with at least 100 assembly barcodes, we found a median of 27.6% perfect assemblies for codon 1 and 22.6% for codon 2, suggesting that the new bead pool can assemble large libraries at high fidelity (Fig. 2b).

DropSynth 2.0 combines improvements in fidelity and scale, significantly enhancing our ability to build large, accurate gene libraries. By improving fidelity, gene libraries enriched with perfect assemblies enable clearer hypothesis testing using multiplexed functional assays. In addition, improvements in fidelity allow for the assembly of longer genes using more oligos. Increasing the frequency of perfect assemblies enables simpler individual gene retrieval using molecular cloning or dial-out PCR^22^. By improving scale, larger gene libraries reduce the cost per assembly reaction and enable more data to be generated on desired hypotheses. Combining these improvements creates a much more powerful workflow for the synthesis of large gene libraries.

## Acknowledgements

This work was supported by the funds from the Human Frontier Science Program [LT000068/2016 to C.P.], Netherlands Organisation for Scientific Research Rubicon fellowship [to C.P.], National Science Foundation Graduate Research Fellowship under Grant No. 2016211460 [to A.M.S], National Institutes of Health New Innovator Award [DP2GM114829 to S.K.], Searle Scholars Program [to S.K.], Department of Energy (DE-FC02-02ER63421 to S.K.), UCLA, and Linda and Fred Wudl. C.P. holds a Career Award at the Scientific Interface from Burroughs-Wellcome Fund.

## Methods

### Oligo Design

The software used to split a given amino acid sequence into oligos with overlaps was derived from Eroshenko et al^18^ and available on https://github.com/KosuriLab/. Amino acid sequences were first converted to nucleic acid sequences by assigning codons randomly weighted based on their frequency in the *E. coli* genome, while also preventing formation of four restriction sites used in cloning and processing (NdeI, KpnI, BtsI-v2, and BspQI). Next, the coding regions were flanked with restriction sites for cloning (NdeI, KpnI) and the forward and reverse assembly primers used in the emulsion polymerase cycling assembly (PCA). The sequences were then split into oligos with overlap regions that satisfy certain parameters, including predicted melting temperature range using the nearest neighbor method^23,24^, mean overlap size, and predicted secondary structure using the hybrid-ss-min function in UNAfold^25^. Sequences that failed to meet these parameters were assigned new codons until a successful split was generated. Split oligo sequences were then flanked with BtsI-v2 sites used to release the oligos inside each droplet. In order to maintain the same length across all oligos, padding sequence consisting of ATGC repeats was added to the region upstream of the 5’ BtsI-v2 site. Next, a Nt.BspQI sequence, 12 nt gene-specific barcode sequence (referred to as the ‘microbead barcode’), and another Nt.BspQI sequence were prepended to the 5’ end of each oligo. Nt.BspQI was used to nick the top strand on the 5’ end of the barcode and the bottom strand on the 3’ end of the barcode sequence, exposing it as a 12 nt top-strand overhang. This barcode allows all oligos contributing to a given gene to be localized on the same bead. Oligos were next flanked with 15 nt amplification primers unique to a given library subpool. BLAT^26^ was run to verify that amplification primer sequences did not possess homologies > 10bp to the designed oligos. Prior to synthesis, final oligo sequences were screened for the presence of all required components and against all illegal restriction sites.

Using the above oligo design, we synthesized a microarray-derived oligo library synthesis (OLS) pools of 33,792 230mer oligos from Agilent Technologies. This pool contained several variations of two codon versions of a 384-member DHFR library derived from our original work^13^. For our control libraries, which were used for all biological optimizations, we used an overlap melting temperature range of 58-62C, mean overlap size of 20bp, and an overlap secondary structure cutoff of −4 kcal/mol. We also generated identical amino acid libraries using alternative overlap parameters, including a longer overlap size of 25bp, and a more stringent secondary structure cutoff of −2 kcal/mol. Another set of amino acid libraries contained alternative IIs restriction sites to BtsI-v2, including BsmAI and BsrDI. This OLS pool also contained 2 codon usages of a single 1536-member DHFR library derived from 4 libraries from Plesa et al^13^.

### Microbead barcode design

In order to generate distinct 12mer barcode sequences, we took 2,000 20mer primer sequences derived from Eroshenko et al^18^, removed all sequences containing NdeI, KpnI, BtsI-v2, BspQI, EcoRI, XhoI, SpeI, and NotI, and generated all possible 12mer subset sequences. We next screened for self-dimers, GC content between 45% and 55% and a melting temperature between 40°C and 42°C, We further filtered sequences to have a minimum modified Levenshtein distance of 3 between selected barcodes^27^. We then selected the first 384 sequences to be used in oligo designs, with complementary sequences being used to generate the beads. For the 1,536-plex barcode design, we performed identical screens except for a relaxed melting temperature screen between 38°C and 44°C. The first 1,536 sequences were used in our 1,536-plex oligo libraries, with complementary sequences being used to generate the beads.

### Barcoded beads protocol

A detailed protocol for barcoded bead preparation is available in the Supplementary Protocols. Three oligos are required to generate each DropSynth barcoded bead, two of which are common to all beads (anchor and ligation oligo). The anchor oligo, which has 5’ double biotin modification, contains sequences complementary to the ligation oligo and part of the barcode oligo. The ligation oligo, which contains 3’ biotin modification and 5’ phosphate modification, is fully complementary to the anchor oligo and allows for the ligation of the barcode oligo. The barcode oligo, which has no modifications, contains a common sequence on the 3’ end which hybridizes to the anchor oligo, and a unique 12nt sequence which acts as a 5’ overhang. This setup minimizes cost, as only the common oligos (anchor and ligation) require expensive modifications. The anchor and ligation oligo were purchased in bulk at >1umole while the barcode oligos were purchased as a single 384-well plate from Integrated DNA Technologies.

The three oligos required for each barcoded bead were individually mixed, ligated, and phosphorylated in individual wells of a 384-well plate using a Liquidator 96 (Mettler-Toledo Rainin, Oakland, CA). Next, magnetic Streptavidin M270 Dynabeads (Invitrogen, Carlsbad, CA) were added to each well, and plates were incubated overnight at room temperature while shaking >2000RPM. The individual wells were then washed >5 times using 2X Bind & Wash Buffer and a 384-Well Post Magnetic Plate (Permagen Labware, Peabody, MA). After washing, individual bound beads were resuspended in 5ul of Bind & Wash Buffer and pooled together. For the 1536-plex barcoded bead pool, 4 plate pools of 384 barcoded beads were combined in equal volumes.

### Oligo amplification and processing

A detailed protocol for oligo amplification, processing, assembly, mismatch binding by MutS, and suppression PCR is available in the Supplementary Protocols. Upon receipt of the oligo pool, individual oligo libraries were PCR-amplified using 15 nt amplification primers with Q5 High-Fidelity 2X Master Mix (New England Biolabs, Ipswitch, MA), and number of cycles determined by qPCR. Amplifications were stopped several cycles prior to plateauing to prevent overamplification. Oligo subpools were then diluted to 0.02 ng/µL and bulk-amplified using a biotinylated forward amplification primer and unmodified reverse amplification primer with Q5 High-Fidelity 2X Master Mix for 20 cycles. For each library, 8 PCRs were run in parallel, pooled and column-cleaned using a Clean & Concentrator (Zymo Research, Irvine, CA). Oligo subpools were then nicked overnight using the nicking endonuclease Nt.BspQI, exposing gene-specific 12 nt barcode overhangs. The short biotinylated fragment cleaved following nicking was removed by binding to Streptavidin M270 Dynabeads (Invitrogen), and the remaining processed oligos were column-cleaned. 1.3 ug of each processed oligo subpool was added to 20ul of barcoded beads (∼5 million beads) and Taq ligase. The mixture was slowly annealed overnight from 50C to 10C, allowing the 12nt overhang on the processed oligos to hybridize to complementary 12nt overhangs on barcoded beads.

### Emulsion assembly

Loaded beads were mixed with a polymerase master mix, either KAPA2G Robust HotStart ReadyMix (KAPA Biosystems, Wilmington, MA), KAPA HiFi HotStart ReadyMix (KAPA Biosystems, or Q5 High-Fidelity 2X Master Mix (New England Biolabs), 60 nt primer sequences containing 20 nt amplification primer sequences and 40 nt ITRs (to be used during bulk suppression PCR), BSA, and BtsI-v2. Immediately after adding BtsI-v2, the mixture was added to 600 µl of BioRad Droplet Generation Oil and vortexed for three minutes using a Vortex Genie 2 (Scientific Industries, Bohemia, NY), resulting in compartmentalization of beads in <5um droplets. After vortexing, samples were aliquoted into PCR strips and incubated at 55°C for 90 minutes, allowing BtsI-v2 to cleave oligo sequences off the beads. Samples were heated to 94°C for 2 min, then thermocycled for 60 cycles with the following conditions: 94°C for 15 sec, 57°C for 20 sec, 72°C for 45 sec, followed by a final 5 min extension at 72°C. Following assembly, emulsions were broken by adding 100µl perfluoro-1-octanol (Sigma Aldrich, Saint Louis, MO), and the aqueous phase was extracted and column-cleaned. Assembled products were then run on a 2% agarose gel and bands were extracted at the correct assembly length.

### Mismatch binding by MutS

Following gel extraction of assembly products, 10µl of M2B2 magnetic beads (US Biological, Salem, MA) was added to each library and incubated for two hours at room temperature while shaking using a Thermomixer C (Eppendorf, Hamburg, Germany). M2B2 beads (US Biological) contain immobilized MutS and thus bind to and magnetically separate DNA containing mismatch-generated heteroduplexes. Following incubation, error-depleted libraries were column-cleaned using a Clean & Concentrator (Zymo Research). In order to verify filtration of DNA, libraries were bulk-amplified on a qPCR using assembly primers before and after M2B2 treatment and ΔCq was quantified.

### Bulk suppression PCR

Gene libraries assembled during DropSynth assembly contain external 40 nt inverted terminal repeats (ITRs) lacking homology to any library sequences. Following recovery of assembled DropSynth libraries, a bulk PCR was carried out using a single 20 nt primer complementary to the proximal region of the 5’ ITR. Due to their close physical proximity, the ITRs of shorter DNA fragments tend to self-anneal, creating hairpin-like structures with suppressed amplification. In contrast, the ITRs of longer DNA fragments are less likely to anneal to one another, allowing for primer annealing and effective amplification. In this case, libraries were amplified using Q5 High-Fidelity 2X Master Mix (New England Biolabs), a final primer concentration of 0.8 µM, Tm of 58°C, and number of cycles determined by qPCR. Amplifications were stopped several cycles prior to plateauing to prevent overamplification. Following amplification, samples were run on a 2% agarose gel and assembly bands were extracted.

### pEVBC plasmid construction

The plasmid used to barcode unique assemblies is derived from our previous work^13^. pEVBC is a pUC19 derivative containing a pLac-UV5 promoter, NdeI and KpnI restriction sites for cloning, an in-frame stop codon and 20mer random assembly barcode sequences. The plasmid was constructed by digesting pUC19 with AatII and BspQI, gel-extracting the larger fragment, and ligating in a gBlock DNA fragment containing the promoter, several restriction sites, and chloramphenicol acetyltransferase in frame before the stop codon. The resulting plasmid was then double digested with NcoI and KpnI and the 2,209bp fragment was gel extracted. Using this fragment as a template, an around-the-horn PCR was carried out using the forward primer pEVBC_FWD containing an NdeI site and reverse primer pEVBC_REV1 containing KpnI and a 20 nt random assembly barcode sequence with the following conditions: 95°C for 3min, followed by 5 cycles of 98°C for 30 sec, 59°C for 15 sec, and 72°C for 3 min. The PCR product was then further amplified using pEVBC_FWD and pEVBC_amp_FWD for 15 cycles. The resulting amplicon was then column purified, digested with NdeI and KpnI, treated with rSAP and size-selected.

### Barcoded library in pEVBC

Following bulk suppression PCR of assembly products, gene libraries were double-digested with NdeI and KpnI and column-purified. Gene libraries were then ligated to digested NdeI + KpnI pEVBC plasmid using a 3:1 insert-to-vector molar ratio, column-purified, and eluted in a volume of 15ul. Ligation products were directly PCR-amplified with sequencing primers mi3_FWD and mi3_N7##_REV to add p5, p7, and indexes (designated by 7##) for Illumina sequencing.

### Assembly barcode sequencing and analysis

Assembly barcoded libraries were sequenced on a total of five Illumina MiSeq paired-end 600-cycle runs. Following PCR amplification with sequencing primers mi3_FWD and mi3_N7##_REV, amplicons were gel-extracted and quantified using an Agilent 2200 TapeStation. Samples were then pooled and sequenced on a MiSeq using custom primers mi3_R1, mi3_R2 and mi3_index, and fastqs were generated for each sample following demultiplexing. In order to eliminate biases in coverage following sequencing, individual fastqs were randomly downsampled to 1,880,288 reads (# of reads of the sample with the lowest read depth). All fastq files were trimmed of adapter sequences with bbduk, and paired-end reads were merged with bbmerge (from BBTools package). Reads were next concatenated and piped into a custom python script, used in our previous work. This script splits reads into variants and 20nt assembly barcodes, generating a dictionary containing each assembly barcode and the variants mapped to it. Assembly barcodes that map to multiple variants were removed by calculating the pairwise Levenshtein distance of every variant associated with a given assembly barcode. If at least 5% of assembly barcodes have a Levenshtein distance > 10, the assembly barcode is considered contaminated and dropped from the analysis. Next, a consensus sequence is generated by taking the majority base call at each position, and translated until the first stop codon. Variants and their mapped barcodes were then imported into R, where they were analyzed for coverage and fidelity. For coverage analyses, the term ‘assemblies represented’ refers to the total number of assemblies corresponding to a perfect amino acid sequence represented by at least one assembly barcode. For fidelity analyses, the term ‘percent perfect assemblies’ is defined as the median percent perfect sequences at the amino acid level determined by using constructs with at least 100 assembly barcodes.

