## Supplementary Information for "DropSynth 2.0: high-fidelity multiplexed gene synthesis in emulsions"

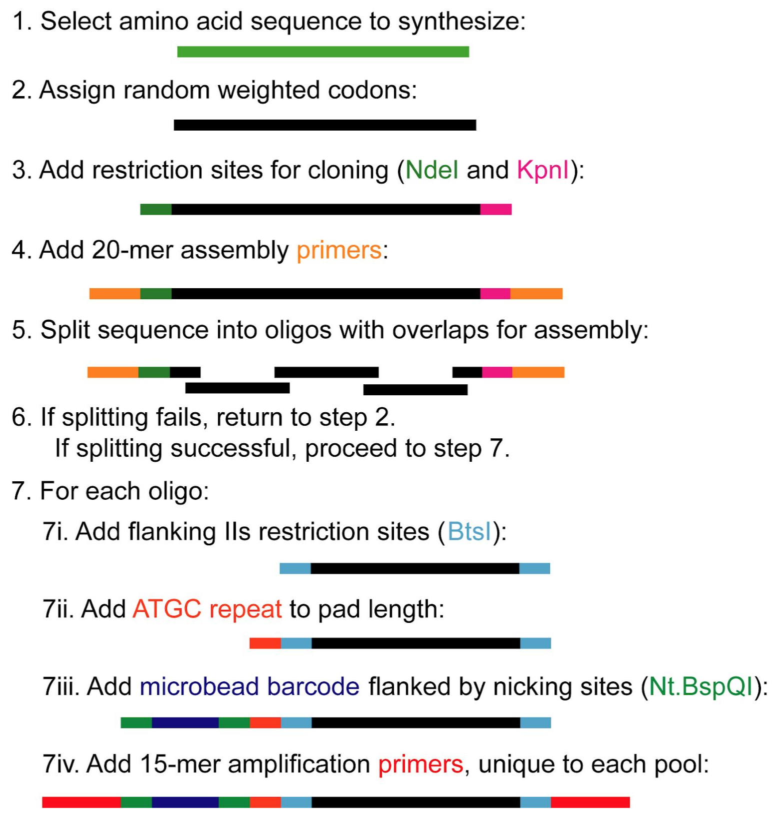


**Supplementary Figure 1**

**Overview of the DropSynth oligo design process.**

The oligo design script, available at https://github.com/KosuriLab/DropSynth and originally derived from Eroshenko et al.[1], takes as input a list of protein sequences and generates all oligos necessary to assemble each gene. First, amino acid sequences are assigned random weighted codons and flanked with restriction sites used for cloning and 20mer assembly primer sequences used for the emulsion assembly. Next, the full gene sequence with restriction sites and primers is split into oligos with overlaps of a predefined length, melting temperature and secondary structure. If splitting fails, which can be due to improper overlap parameters, long homopolymers, or forbidden restriction sites, the protein sequence is reassigned new random weighted codons and the process is repeated. Once each gene is successfully split into oligos, each oligo is flanked with BtsI sites used to cleave sequences off beads, padding sequence, a 12mer gene-specific microbead barcode sequence flanked by Nt.BspQI sites, and 15mer amplification primer sequences used to amplify the oligo libraries from the OLS pool.

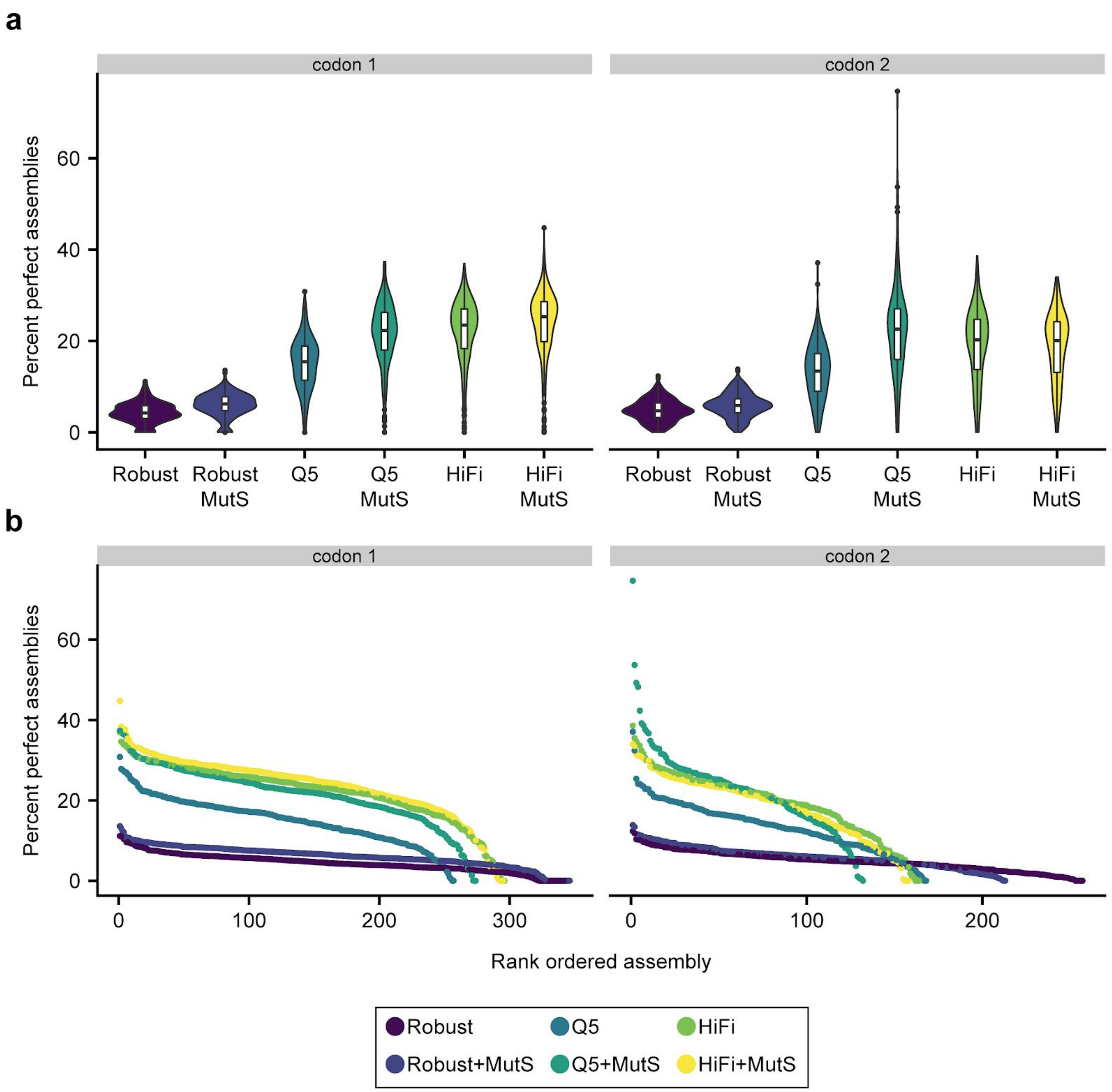


**Supplementary Figure 2**

**DropSynth assembly of 2 codon versions of a 384-gene library using 3 different polymerases with or without MutS-based enzymatic error correction.**

**a,** Comparison of percent perfect assemblies (minimum 100 assembly barcodes) of 2 codon versions of a 384-gene library assembled using DropSynth with 3 different polymerases (KAPA Robust, NEB Q5, or KAPA HiFi) with or without MutS-based enzymatic error correction. **b,** Rank ordered plot of percent perfect assemblies (minimum 100 assembly barcodes) of all conditions. Though assemblies with KAPA Robust have the greatest library representation, assemblies with high-fidelity polymerases NEB Q5 and KAPA HiFi have significantly improved fidelity of represented constructs.

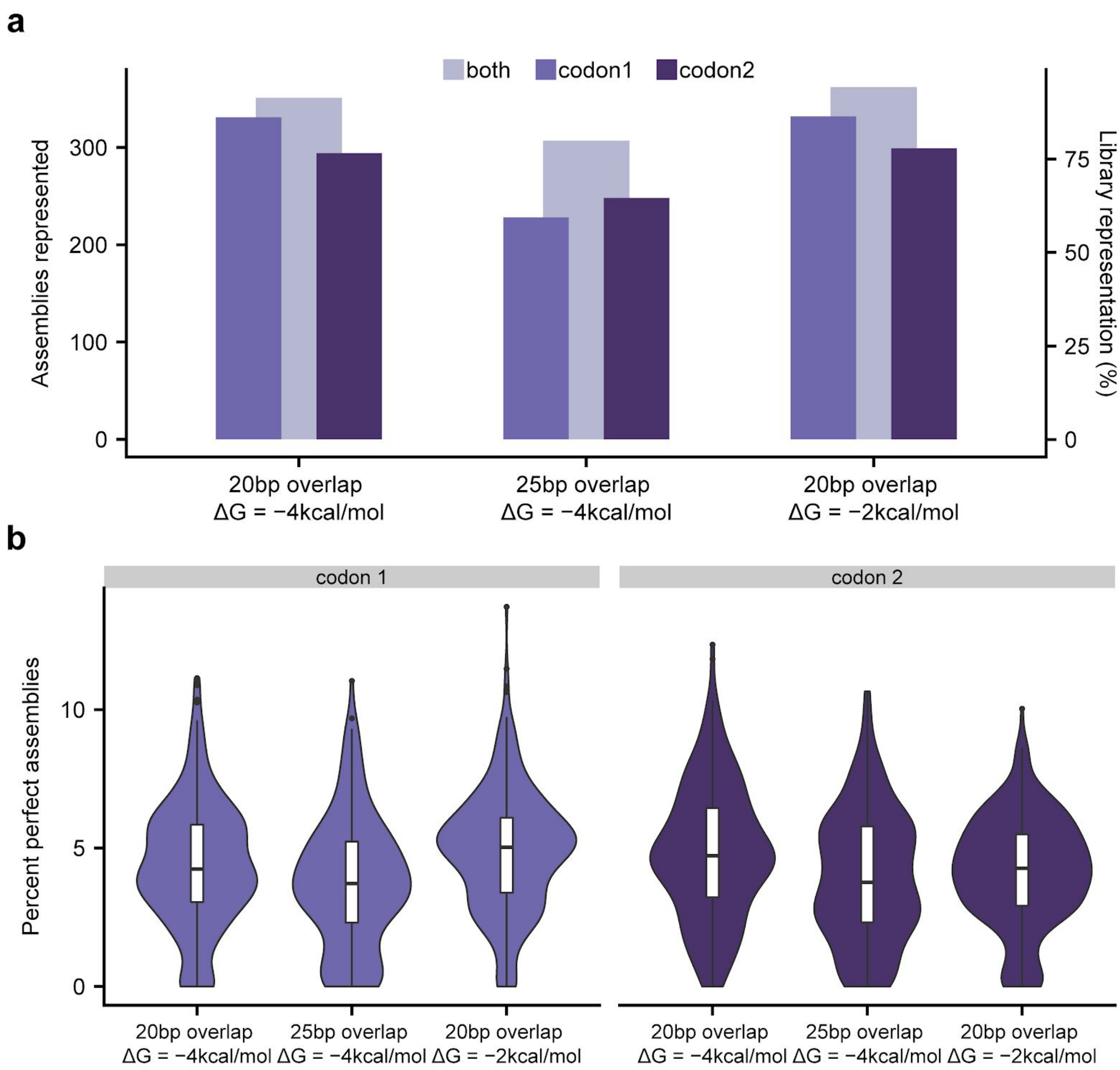


**Supplementary Figure 3**

**DropSynth assembly of 2 codon versions of a 384-gene library containing alternative oligo overlap parameters (length, secondary structure).**

**a,** Comparison of total assemblies represented with at least one assembly barcode of 2 codon versions of a 384-gene library designed with alternative average overlap lengths (20 or 25bp) and overlap secondary structure thresholds (maximum ΔG = -4 kcal/mol or -2 kcal/mol) and assembled using DropSynth with KAPA Robust. Modifying the overlap secondary structure appears to have little effect on representation, while increasing the average overlap length to 25bp has a slight negative effect on representation. **b,** Comparison of percent perfect assemblies (minimum 100 assembly barcodes) of all conditions.

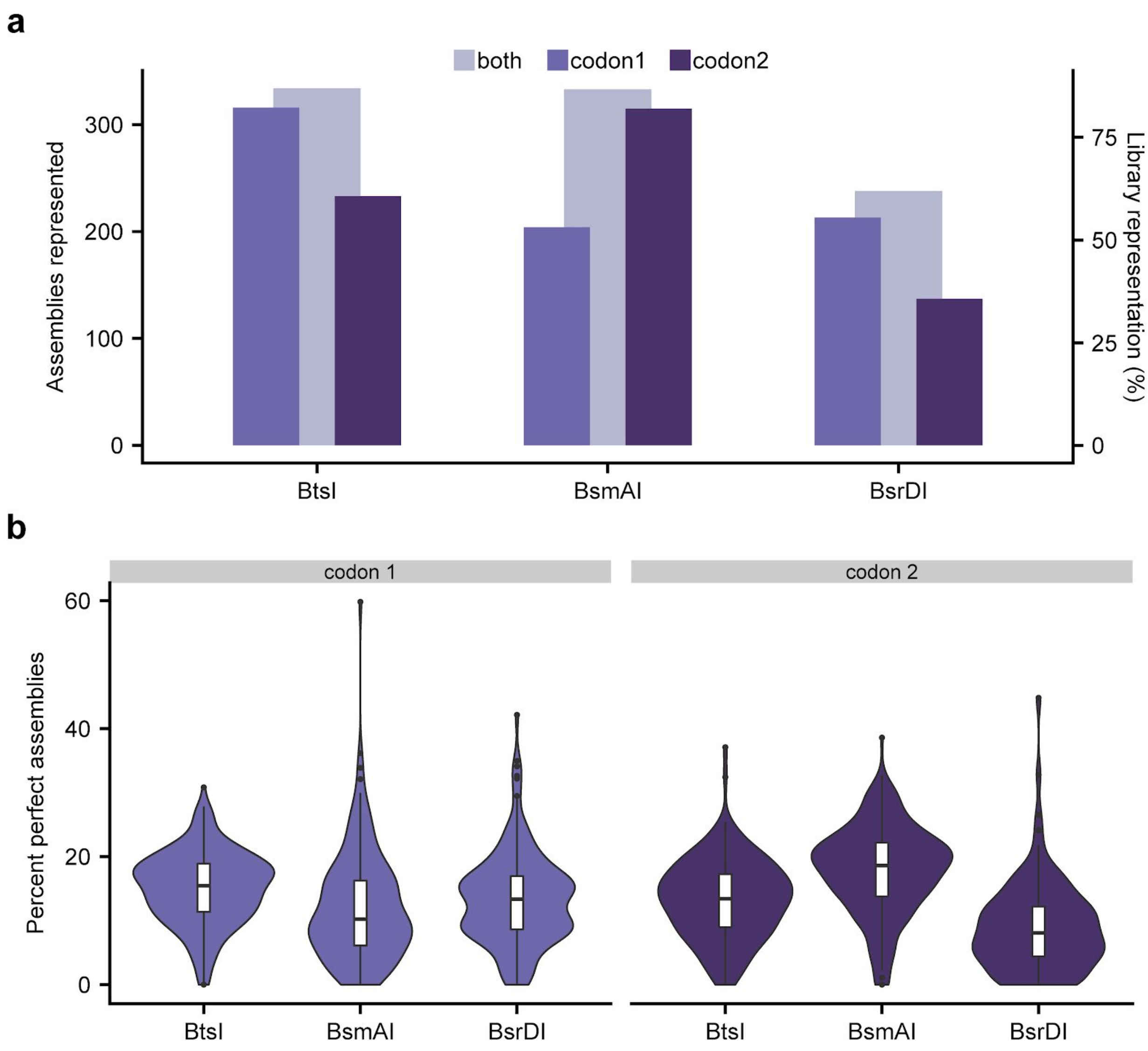


**Supplementary Figure 4**

**DropSynth assembly of 2 codon versions of a 384-gene library containing alternative IIS restriction sites (BtsI, BsmAI, and BsrDI).**

**a,** Comparison of total assemblies represented with at least one assembly barcode of 2 codon versions of a 384-gene library designed with alternative IIS restriction sites used to cleave oligos off the beads (BtsI, BsmAI, or BsrDI) and assembled using DropSynth with NEB Q5. Using BsrDI appears to have a slight negative effect on representation compared to BtsI and BsmAI. **b,** Comparison of percent perfect assemblies (minimum 100 assembly barcodes) of all conditions.

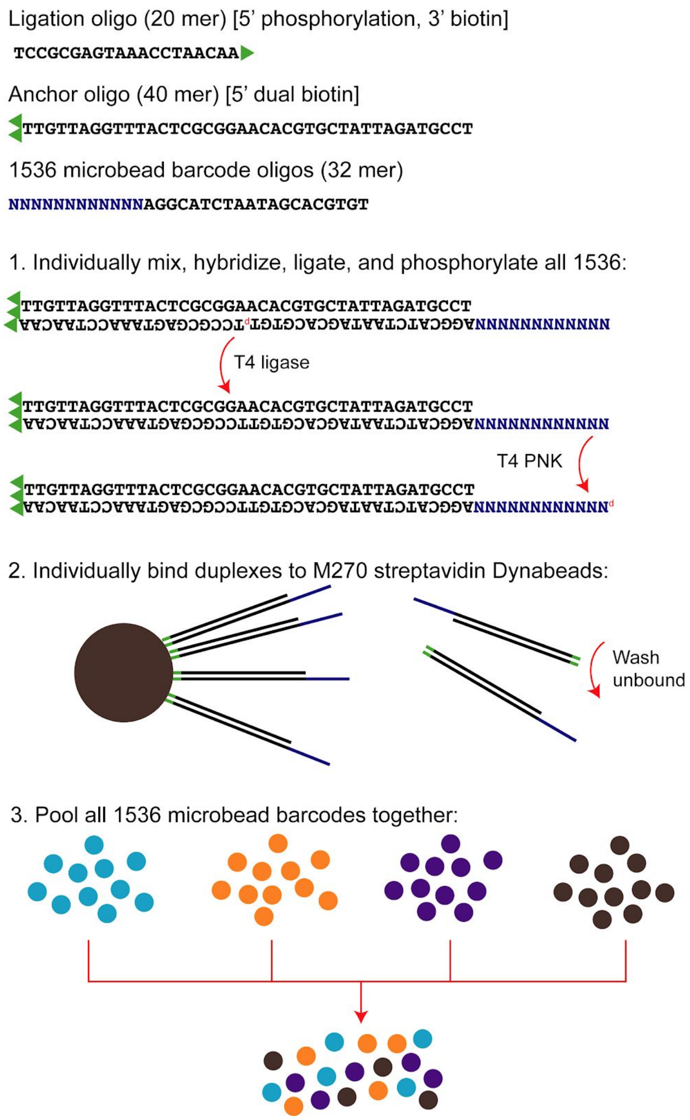


**Supplementary Figure 5**

**Overview of the 1536-plex barcoded bead generation process.**

The 1536-plex barcoded bead generation process is derived from the 384-plex bead generation process originally demonstrated in Plesa et. al[2]. The process requires 3 oligos: a 20mer ligation oligo with 5' phosphorylation and 3' biotinylation, a 40mer anchor oligo with 5' dual biotinylation, and 1536 32mer microbead barcoded oligos. Each microbead barcoded oligo is individually hybridized to the anchor and ligation oligos in 4 384-well plates, forming three-oligo complexes with 12nt 5' overhangs containing the designed 12mer microbead barcode sequences. T4 ligase then seals the nick between the ligation and microbead barcoded oligo, and T4 PNK phosphorylates the 12nt 5'microbead barcode overhang. All duplexes are then individually bound to M270 Streptavidin Dynabeads, washed, and pooled to form a single 1536-plex barcoded bead pool.

**Supplementary Protocols**

**DropSynth 2.0 bead barcoding protocol**

This protocol can be performed using 1 384-well plate to generate 384 unique barcoded beads, or 4 384-well plates to generate 1536 unique barcoded beads. Though the process can be done by hand, it is helpful to use a Rainin Liquidator 96 for liquid handling steps.

Reagents Required (384-plex):

1. 240 µL 100 µM anchor oligo (Integrated DNA Technologies)
2. 240 µL 100 µM ligation oligo (Integrated DNA Technologies)
3. 1,056 µL 10X T4 Ligase Buffer (New England Biolabs)
4. 1 µL of each 100 µM barcoded oligo (Integrated DNA Technologies)
5. 24 µL T4 Ligase (New England Biolabs)
6. 240 µL T4 PNK (New England Biolabs)
7. 1500 µL Streptavidin M270 Dynabeads (Invitrogen)
8. >10 mL UltraPure Distilled Water (Invitrogen)
9. >10 mL 2X B&W Buffer

Reagents Required (1536-plex):

1. 960 µL 100 µM anchor oligo (Integrated DNA Technologies)
2. 960 µL 100 µM ligation oligo (Integrated DNA Technologies)
3. 4,224 µL 10X T4 Ligase Buffer (New England Biolabs)
4. 1 µL of each 100 µM barcoded oligo (Integrated DNA Technologies)
5. 96 µL T4 Ligase (New England Biolabs)
6. 960 µL T4 PNK (New England Biolabs)
7. 6,000 µL Streptavidin M270 Dynabeads (Invitrogen)
8. >40 mL UltraPure Distilled Water (Invitrogen)
9. >40 mL 2X B&W Buffer

Prepare 40mL 2X B&W buffer (2M NaCl, 1mM EDTA, 10mM Tris):

- 4.675g NaCl salt
- 400 µL UltraPure 1M Tris, pH 7.5 (Invitrogen)
- 80 µL UltraPure 0.5 M EDTA, pH 8.0 (Invitrogen)
- UltraPure Distilled Water (Invitrogen) to 40 mL

1. Hybridize the anchor, ligation and barcoded oligos:
    - Add to the first row of 96-well deep well plate:
        - 20 µL 100 µM anchor oligo
        - 20 µL 100 µM ligation oligo
        - 80 µL 10X T4 Ligase Buffer
        - 640 µL UltraPure Distilled Water
    - Using a Rainin P200 12-channel pipette, add 95 µL of master mix to all rows of master 96-well plate.
    - Using a Rainin Liquidator 96, distribute 19 µL of master mix from master 96-well plate to all wells of a new 384-well plate. The 384-well plate can be adjusted to 4

    corners using a Rainin Plate Adapter 384, allowing all wells to be filled from the 96-well master plate.
    - Using a Rainin Liquidator 96, transfer 1 µL from every well of the 100 uM barcoded oligo plate to every well of the 384-well plate.
    - Anneal the mixed oligos on each plate using the following conditions:
        - 3 min at 70ºC
        - Ramp down to 60ºC for 1 min, 0.1ºC/sec
        - Ramp down to 50ºC for 1 min, 0.1ºC/sec
        - Ramp down to 40ºC for 1 min, 0.1ºC/sec
        - Ramp down to 30ºC for 1 min, 0.1ºC/sec
        - Put plate on ice
2. Ligate the barcoded oligo to the ligation oligo:
    - Add to the first row of a 96-well plate:
        - 2 µL T4 Ligase
        - 8 µL 10X T4 Ligase Buffer
        - 70 µL UltraPure Distilled Water
    - Using a Rainin P20 12-channel pipette, add 10 µL of master mix to all rows of a master 96-well plate.
    - Using a Rainin Liquidator 96, distribute 2 µL master mix from the master 96-well plate to all wells of the 384-well plate.
    - Incubate plate at 16ºC for 1 hr or longer, followed by 65ºC for 20 min to heat inactivate the ligase.
3. Phosphorylate the barcoded oligo:
    - Add to first row of 96-well plate:
        - 20 µL T4 PNK
        - 60 µL UltraPure Distilled Water
    - Using a Rainin P20 12-channel pipette, add 10 µL of master mix to all rows of a master 96-well plate.
    - Using a Rainin Liquidator 96, distribute 2 µL master mix from the master 96-well plate to all wells of the 384-well plate.
    - Incubate the plate at 37ºC for 40 min (or longer), followed by 65ºC for 20 min to heat inactivate the PNK.
4. Bind to beads:
    - Prepare 1500 µL stock Dynabeads M270 Streptavidin, washed, and resuspended in 3000 µL 2X B&W buffer.
    - Add 200 µL to first row of 96-well plate.
    - Using a Rainin P200 12-channel pipette, add 25 µL of master mix to all rows of a master 96-well plate.
    - Using a Rainin Liquidator 96, add 5 µL resuspended beads to each well of the 384-well plate.
    - Mix overnight with shaking (>2000 RPM) at room temperature.
5. Pool beads:
    - Using a Rainin Liquidator 96, wash each well with 20 µL 2X B&W buffer 8 times.
    - Using a Rainin Liquidator 96, resuspend each well in 5 µL 2X B&W buffer.
    - Mix 5 µL of each well together, making a 1920 µL mixed barcoded bead pool for each plate. Store these at 4ºC when not in use.

**DropSynth 2.0 emulsion synthesis protocol**

1. Prepare the OLS pool
    - Make a 1/10 dilution of the OLS chip pool.
    - Prepare mixtures of forward and reverse subpool amplification primers for each subpool, with 10 µM final concentration of each primer.
2. Amplify subpools.
    - For each subpool, run a qPCR to determine the number of cycles required for amplification. Amplifications are stopped several cycles before plateauing to prevent over-amplification of the libraries.
    - Amplify each subpool using NEB Q5.
        - 1 µL template (1/10 OLS pool dilution)
        - 1.25 µL subpool specific primer 10 µM ampF
        - 1.25 µL subpool specific primer 10 µM ampR
        - 21.5 µL UltraPure Distilled Water (Invitrogen)
        - 25 µL NEBNext Q5 Hot Start HiFi PCR Master Mix (New England Biolabs)
        - TOTAL: 50 µL
        - PCR protocol:
            1. 45 sec 98ºC initial denaturation
            2. 15 sec 98ºC denaturation
            3. 30 sec 58ºC annealing
            4. 15 sec 72ºC extension
            5. Go to step 2, repeat based on the number of cycles determined by qPCR.
            6. 1 min 72ºC final extension
    - Column purify amplified oligos using a DNA Clean & Concentrator -5 (Zymo Research).
    - Run PCR products on gel. Look for higher MW products, indicative of overamplification. Excessive low MW products may indicate chip synthesis issues.
    - Size select, using gel extraction, if necessary.
    - Create 20 pg/µL dilutions of each amplified subpool.
3. Bulk amplify subpools.
    - Run a second PCR using a <u>biotinylated</u> FWD amplification primer, with sufficient tubes to make 5 ug to 10 ug of PCR product.
        - 1 µL of 20 pg/µL subpool dilution
        - 1.25 µL subpool specific primer mix 10 µM biotinylated ampF
        - 1.25 µL subpool specific primer mix 10 µM biotinylated ampR
        - 21.5 µL UltraPure Distilled Water (Invitrogen)
        - 25 µL NEBNext Q5 Hot Start HiFi PCR Master Mix (New England Biolabs)
        - TOTAL: 50 µL

        - PCR protocol:
            1. 45 sec 98ºC initial denaturation
            2. 15 sec 98ºC denaturation
            3. 30 sec 58ºC annealing
            4. 15 sec 72ºC extension
            5. Go to step 2, 18X
            6. 1 min 72ºC final extension
    - Pool and column purify using a DNA Clean & Concentrator -25 (Zymo Research).
4. Nicking.
    - Nick the bulk amplified subpools. Split the following across multiple tubes depending on the amount of DNA to be processed. In each 1.5 mL tube add:
        - 15 µL Nt.BspQI (10U/µL) (New England Biolabs)
        - 5 to 10 ug of DNA
        - 15 µL NEBuffer3.1 (New England Biolabs)
        - UltraPure Distilled Water (Invitrogen) to 150 µL total
    - Leave at 50ºC overnight with shaking >1500 RPM.
5. Capture and remove the short biotinylated fragment.
    - Wash 50 µL streptavidin M270 Dynabeads (Invitrogen) for each 1.5 mL tube in the nicking reaction, as per manufacturer's instructions and resuspend in 2X B&W buffer.
    - Add 50 µL of washed beads to the 150 µL nicking reaction in each tube.
    - Incubate at 55ºC with 800 RPM shaking for at least 1 hour.
    - Move all 1.5 mL tubes to a 55ºC water bath.
    - Place the tube so that solution is just below the surface of the water. Hold a strong magnet underwater against the side of the tube to magnetically separate Dynabeads. Pipette the supernatant, which contains the processed oligos and save them in a new container. Remove the tube with the Dynabeads from the magnet.
    - Add 100 µL of UltraPure Distilled Water (Invitrogen) to the tube and resuspend the beads. Incubate these at 55ºC for another 30 min and then repeat the procedure to recover the supernatant again while leaving the Dynabeads behind.
    - Repeat this procedure for all tubes as necessary.
    - Pool processed oligos (supernatant) for each subpool and column purify using a DNA Clean & Concentrator -5 (Zymo Research).
6. Capture processed oligos with barcoded beads.
    - Take 20 µL of the pooled barcoded beads. These are in stored in 2X B&W buffer (high ionic concentration) which may interfere with ligation reaction. Resuspend them in 20 µL UltraPure Distilled Water (Invitrogen).
    - Mix the processed DNA with the barcoded beads:
        - 1.3 ug processed DNA (~12 pmol)
        - 20 µL pooled barcoded beads (~6 million beads, binding capacity 1.3 ug DNA)
        - 10 µL 10X Taq ligase buffer (New England Biolabs)

        - 4 µL Taq ligase (40 U/µL) (New England Biolabs)
        - UltraPure Distilled Water (Invitrogen) to 100 µL
    - Overnight cycling (>2 hr incubation at each of the following temperatures) (13 hr), use shaking to prevent beads from settling down:
        - 3 hours @ 50ºC
        - Ramp to 40ºC for 3h, 0.1ºC/sec
        - Ramp to 30ºC for 3h, 0.1ºC/sec
        - Ramp to 20ºC for 2h, 0.1ºC/sec
        - Ramp to 10ºC for 2h, 0.1ºC/sec
    - Wash 3 times at 4ºC using 2X B&W buffer. This is important for removing unbound oligos in order to increase specificity.
    - Wash twice at RT using 2X B&W buffer
    - Re-suspend in 100 µL Elution Buffer (Qiagen) (~60k beads/µL)
7. Emulsion assembly (ePCA).
    - Setup emulsion. All of this procedure should be done on ice. FWD and REV assembly primers contain ITR overhangs which will be used for single-primer suppression PCR.  Add BtsI-v2 only at the very last step. Try to minimize the time between adding the BtsI-v2 and vortexing the emulsion.
        - 40 µL of loaded beads (~500 ng DNA)
        - 0.5 µL 100 µM AsmF_40bpITR
        - 0.5 µL 100 µM AsmR_40bpITR
        - 50 µL KAPA HiFi 2X Mastermix (KAPA Biosystems)
        - 1 µL BSA (New England Biolabs)
        - 1 µL UltraPure Distilled Water (Invitrogen)
        - 7 µL BtsI-v2 (New England Biolabs) (add last)
        - TOTAL: 100 µL
    - Mix at low speed in vortexer to resuspend beads.
    - Add 600 µL Droplet Generation Oil for EvaGreen (Bio-Rad) to a 1.5mL non-stick tube.
    - Add 100 µL aqueous phase to the bottom of the oil phase.
    - Vortex at Max Speed in foam holder taped down for 3 minutes. If doing multiple emulsions, do this one at a time. We use a Vortex Genie 2 (Scientific Industries) at max speed.
    - After vortexing all emulsions, place each emulsion into PCR tubes with 100 µL in each tube. Use a P1000 tip to avoid disturbing the emulsion. Most of the droplets will float to the top of the tube, try to get as much of this as possible and distribute this over multiple PCR tubes.
    - PCR Cycling
        - 55ºC for 90 min (allow BtsI-v2 to cleave DNA from the beads)
        - 94ºC for 2 min (initial denaturing)
        - 94ºC for 15 sec (denaturing)
        - 57ºC for 20 sec (annealing)
        - 72ºC for 45 sec (extension)

        - Go to step 3 for additional 60 cycles
        - 72ºC for 5min (final extension)
        - 4ºC forever
8. Break the emulsion:
    - After ePCA, pipet out the entire volume of droplets from each PCR tube into a 1.5 mL tube.
    - Add 100 µL of 1*H*,1*H*,2*H*,2*H*-Perfluoro-1-octanol (Sigma Aldrich) for each 100 µL of PCR reaction combined in the 1.5mL tube.
    - Vortex at maximum speed for 1 min.
    - In a centrifuge, spin down at 15,500 x g for 10 min.
    - If droplets are still present, vortex and centrifuge again.
    - Remove upper aqueous phase by pipetting, avoiding the oil phase.
    - Transfer this to a clean 1.5mL tube (this is the DNA).
    - Column purify using a DNA Clean & Concentrator -5 (Zymo Research).
9. Size selection.
    - The amplicons will often be mixed with undesired lower-molecular weight assemblies. Removing these using size selection will increase final yield.
        - Gel extraction
            1. Run amplicons on a gel and extract the correct range and purify.
            2. Note: Typically there is not enough DNA after the ePCA to visualize on a gel, so this is often a blind extraction.
10. MutS treatment (optional)
    - Enzymatic error correction can be used to enrich for perfect assembly products. Here we use M2B2 magnetic beads (US Biological), which contain immobilized MutS and thus bind to and magnetically separate DNA containing mismatch-generated heteroduplexes.
    - Add 10 µL of M2B2 magnetic beads to size-selected assembly product.
    - Incubate at 20ºC with 1600 RPM shaking for at least 1 hour.
    - Immediately place on magnetic rack and extract supernatant.
    - Column clean the DNA using a DNA Clean & Concentrator -5 (Zymo Research).
11. Single-primer suppression PCR.
    - In this technique, self-annealing of inverted terminal repeats (ITRs) flanking the assembled genes competes with the annealing of a single primer which aligns to part of the ITR[3]. Shorter by-products tend to self-anneal, while correct assembly products anneal to the primer, resulting in proper amplification.
        - 1 µL template
        - 4 µL 10 µM suppression primer
        - 25 µL NEBNext Q5 Hot Start HiFi PCR Master Mix (New England Biolabs)
        - UltraPure Distilled Water (Invitrogen) to 50 µL
        - PCR protocol:
            1. 3 min 95ºC initial denaturation
            2. 15 sec 98ºC denaturation

3. 30 sec 58ºC annealing
4. 15 sec 72ºC extension
5. Go to step 2, determine cycles using qPCR.
6. 1 min 72ºC final extension

- Column purify using a DNA Clean & Concentrator -5 (Zymo Research).
- Check size distribution on gel or tapestation.
- Quantify DNA and proceed to downstream applications.

**Supplementary Table 1**: The oligos required for the bead barcoding process. All oligos were ordered from Integrated DNA Technologies.

| Oligo Name | Sequence | Modifications |
|---|---|---|
| Ligation oligo | TCCGCGAGTAAACCTAACAA | 3' biotin<br>5' phosphorylation |
| Anchor oligo | TTGTTAGGTTTACTCGCGGAACACGTGCTATTAGATGCCT | 5' dual biotin |
| Barcoded oligos | 12-mer DropSynth barcode reverse complement + AGGCATCTAATAGCACGTGT | none |

**Supplementary Table 2**: The oligos required for ePCA and single-primer suppression PCR.The suppression primer aligns to the proximal 20bp of the ITR overhang. All oligos were ordered from Integrated DNA Technologies.

| Oligo Name | Sequence |
|---|---|
| AsmF_40bpITR | TAAGCGCCCTTCTAATACCCAGGTCTGGCCCTATATACGAATCGGGGATGGTAACTAACG |
| AsmR_40bpITR | TAAGCGCCCTTCTAATACCCAGGTCTGGCCCTATATACGAATAGCTGATTGTCCGTTGGT |
| Suppression primer | AGGTCTGGCCCTATATACGA |

**Supplementary Table 3**: The primers required to amplify libraries from the OLS pool. All oligos were ordered from Integrated DNA Technologies.

| Lib | Codon | AmpF Name | AmpF Sequence | AmpR Name | AmpR Sequence |
|---|---|---|---|---|---|
| Control | 1 | skpp15-9-F | CGATCGTGCCCACCT | skpp15-9-R | GTGCGGGCTCCAACT |
| Control | 2 | skpp15-13-F | GGGTTCGAGCGGGAG | skpp15-13-R | TAGCGCGCAGAGAGG |
| Overlap | 1 | skpp15-23-F | AGCTGCTACACCGCC | skpp15-23-R | GCGCGATGGTCACAG |
| Overlap | 2 | skpp15-26-F | GCGGCACCACAAACT | skpp15-26-R | CGTGGCCTCTGTCCT |
| ∆G | 1 | skpp15-30-F | TCCACCGTCGGCAAG | skpp15-30-R | GGCCGCACCCAGTAG |
| ∆G | 2 | skpp15-33-F | AAGTGCCCTTCCCGT | skpp15-33-R | GAGTCCGCGCAAGAG |
| BsmAI | 1 | skpp15-40-F | AGGCGGTCGAGAGTG | skpp15-40-R | CCGTCCTCCACCCAG |
| BsmAI | 2 | skpp15-46-F | CCGCATGCAGTCCCT | skpp15-46-R | CGACTCTTGCGCCCT |

| | | | | | |
|---|---|---|---|---|---|
| BsrDI | 1 | skpp15-49-F | GGCCCAGCGAAGATG | skpp15-49-R | GATCAGCACCGCGAC |
| BsrDI | 2 | skpp15-51-F | GGCGCGCTCTAACAC | skpp15-51-R | CTCCCTCTCGCAGCA |
| 1536 | 1 | skpp15-56-F | AACGCCCAGCCTGTC | skpp15-56-R | CCGCGTTGCTGAGTG |
| 1536 | 2 | skpp15-59-F | AGGCACGCTCAACCT | skpp15-59-R | CCTAGGTCGCACGCA |
